# A functional annotation of a selected protein-coding gene in the *Olimarabadopsis pumila* genome: Oxysterol-binding protein-related protein (ORP) 2B

**DOI:** 10.1101/2025.08.13.670020

**Authors:** Noah Rubin

**Affiliations:** Department of Molecular, Cell, and Developmental Biology, University of California, Los Angeles, Los Angeles, California, United States of America

## Abstract

*Olimarabidopsis pumila*, a spring ephemeral from Asia, is closely related to the well-studied model organism *Arabidopsis thaliana*, though with various important differences. *O. pumila* has evolved to possess various adaptations to its arid environment, including a high salt-tolerance and utilization of extreme light conditions, which *A. thaliana* does not have. However, *O. pumila* is drastically understudied, despite the advantages of using natural genetic variation and broader contexts in researching the various fields of genomic, cellular, and plant biology. In this study, we attempted to address this lack of important research by contributing to the functional annotation of the genome of *O. pumila*. One gene was selected and analyzed using various genomic biology softwares — particularly GenSAS, AUGUSTUS, NCBI BLASTP and MSA Viewer, PSIPRED, AlphaFold, InterPro, STRING, DeepTMHMM, SignalP, TargetP, and WoLF PSORT — to determine its sequence conservation, three-dimensional structure, function, and subcellular localization. The gene, located on contig tig00000022 of the *O. pumila* genome, was found to encode oxysterol-binding protein-related protein (ORP) 2B. We concluded this protein is involved in sterol transportation from the endoplasmic reticulum, and possibly works in contribution to the important biological processes of cell proliferation and division.

## Introduction

The plant *Arabidopsis thaliana* (*A. thaliana*), of the mustard family *Brassicaceae*, has been used as a model organism in several fields of scientific research, from systems biology[1] to genome analysis[2]. The species and its genome has been widely studied, a fact demonstrated by its being the first flowering plant to have its entire genome sequenced about 25 years ago[3].

While *A. thaliana* is a useful tool as a model organism, it is important to expand study to other plants, such as other plants in the *Brassicaceae* family. Differences among species enable researchers to investigate key genes across diverse genetic backgrounds, thereby broadening insights beyond a single model system. In studying *A. thaliana*’s relative species, researchers can apply what has been learned by studying the model organism to new, yet similar, species. This natural source of genetic variation also enables studying of the genes without needing laboratory-induced mutants[4].

However, related species of *A. thaliana* and their genomes are not well studied, particularly *Olimarabidopsis pumila* (*O. pumila*; also sometimes called *Arabidopsis pumila*). *O. pumila* is an ephemeral plant from the arid regions of central, south, and southwest Asia that is very closely related to *A. thaliana. O. pumila*’s genome reflects this relation, with genetic similarities to *A. thaliana*, including transposable elements like Tag1[5]. However, *O. pumila* does also possess many differences in its genetic code, such as in certain rapidly evolving reproductive genes[6] and in its unique tolerance for high salinity[7]. These adaptations not only differentiate *O. pumila* from *A. thaliana*, but also align it with more distantly related and understudied organisms, such as in its shared high-light adaptations with *Sisymbrium altissimum[8]*. Thus, while *O. pumila* and *A. thaliana* share many genomic features, the unique genetic differences in *O. pumila* make it a valuable model for a wide range of biological studies. To date, however, only 206 of its proteins have been characterized, compared with over 130,000 annotated genes in *A. thaliana[9]*, highlighting the urgent need for more comprehensive research.

In this study, we used various gene annotation tools to understand a selected gene from *O. pumila*’s sequenced genome in order to contribute to the functional genomic research of this species. Genes surveyed were located between base pairs 16,500,000 and 17,000,000 on contig tig00000022 and were chosen primarily for their similarity to homologous genes in related species, as determined by BLAST alignments. The ultimately selected gene was located at base pairs 16,594,582-16,599,948, and was found to encode for the oxysterol-binding protein-related protein 2B. This protein is hypothesized to be involved in sterol transportation, particularly to or from the endoplasmic reticulum (ER), and the biological process that it contributes to may be important for cell proliferation or division.

## Materials and Methods

### Genome sequencing and assembly

A previously sequenced and assembled *O. pumila* genome from sequencing data collected by researchers at the University of California, Los Angeles (UCLA) was used.

### *Ab initio* gene prediction

The genome sequences were previously uploaded onto the Genome Sequence Annotation Server (GeneSAS, located at https://www.gensas.org/), a web-based platform for structural and functional genome annotation using an integrated JBrowse and Apollo browser[10]. In GenSAS, the gene prediction tool AUGUSTUS was used.

AUGUSTUS completes *ab initio* gene prediction by locating and labeling structural elements such as transcription start sites, coding and non-coding regions, and splice sites[11]. For this paper, predicted genes were chosen from the contig tig00000022 in the base pair region 16,500,000-17,000,000 of the *O. pumila* genome, with emphasis on proteins between 250 and 750 amino acid residues in length as predicted by AUGUSTUS.

### Protein alignment

The National Center for Biotechnology Information (NCBI) Protein Basic Local Alignment Search Tool (BLASTP, found at https://blast.ncbi.nlm.nih.gov/Blast.cgi) was used to assess the alignment of the AUGUSTUS-predicted protein sequence with a vast database of proteins, including calculating the statistical significance of similarities[12]. The provided FASTA sequence from AUGUSTUS of the selected proteins were inputted into the query search of BLASTP with all default settings, generating outputs that included the most similar homologs across the protein database and their percent identity to and percent coverage of the query sequence, as well as an E-value that showed statistical probability of chance of similarity and an accession code for information of the homolog. To observe conservation and differences between the query *O. pumila* protein and its homologs, one aligned protein was chosen from each species, and these were chosen based on high percent query coverage and high percent identity (above 85% for each), an E-value on a lower order of magnitude than 10^−5^, and with priority for accession codes beginning with NP or XP, which indicate reliably curated genes and computationally predicted genes, respectively. These parameters were set so as to avoid analyzing duplicate sequences from within a species, and to ensure that the aligned homologs are as similar to the query protein as possible to assist functional analysis. For each query, approximately 7-9 homologs were chosen to assess conservation.

After homologs were selected, the Multiple Sequence Alignment (MSA) Viewer (version 1.26.0) built into the NCBI BLAST website was used to visualize similarity between the query protein and selected homologs. MSA Viewer shows where the protein sequences are the same and where they differ, including both small point differences — that may have arisen due to evolutionary changes, individual point mutations, or sequencing errors — and larger differences like insertions and deletions, as well as segments that are largely variable between homologs. Similarities are shown in red, while different sequences are shown in gray, and variable segments are blue. Homologs with insertions compared to the query are shown with a blue bracket at the site of distinction, and deletions relative to the query are indicated by a thin red line[13]. The overall amount of similarity between homologs and query proteins — especially regarding insertions, deletions, and variable regions — were used to further narrow down *O. pumila* genes of interest, preferring fewer major differences relative to other queries.

### Structural analysis

The secondary structure of peptide sequences of interest was visualized by the Position-Specific Iterated BLAST-based Secondary Structure Prediction (PSIPRED) tool (version 4.0, located at https://bioinf.cs.ucl.ac.uk/psipred/)[14]. The protein sequence was pasted into the query search with default settings, and a visualization of amino acid residues involved in various local secondary structures was outputted.

The AlphaFold 3 server (https://alphafoldserver.com/) visualized the three-dimensional structure of proteins of interest using an artificial intelligence-based model. AlphaFold allows researchers to visualize either a single peptide of the inputted sequence, or multiple copies of the same peptide in order to see if the overall protein is composed of multiple subunits of the query peptide. The output of a query generates several predicted accuracy scores, including the plDDT (predicted Local Distance Difference Test) score that measures the confidence of a single amino acid residue’s predicted position and structural participation, and the pTM (predicted Template Modeling) score that measures the overall confidence of the protein structure. A pTM score of more than 0.5 indicates that the predicted structure may be similar to the query protein’s true natural structure[15]. If the pTM score decreased with increasing copies of the peptide, then it was assumed that the protein consisted of only one subunit of the encoded peptide; if the pTM score increased with increasing copies, then the opposite was assumed and the query copies settings with the highest pTM score was assumed to be the most likely natural number of subunits.

### Functional analysis

InterPro (https://www.ebi.ac.uk/interpro/)was used to help determine protein function through the identification of conserved functional domains within query sequences. The resource works by integrating predictive models from various databases to classify sequences according to protein families and predict the presence of specific domains or other significant sites[16]. The FASTA sequence of query proteins was pasted into the search box and default settings options were used. InterPro maps conserved domains and protein families onto the sequence, as well as predicted regions of intrinsic disorder. The output also included potential gene ontology (GO) terms regarding the protein’s biological process, molecular function, and cellular component.

The Search Tool for the Retrieval of Interacting Genes/Proteins (STRING) database (version 12.0, located at https://www.string-db.org/) was used for further functional analysis by indicating how homologs interact directly or indirectly with other proteins, as evidenced by genomic context predictions, high-throughput experiments, conserved co-expression, previous knowledge published in databases, and automated textmining[17]. In this study the FASTA sequence was inputted in the “Proteins by sequences” search box, and the organism was set as the well-studied model organism *A. thaliana*. The results of the search include a list of various proteins from *A. thaliana* that match the inputted sequence to varying degrees, as indicated by a percent identity and an E-value, similar to the E-value in BLAST (see above). For this paper, the protein with the highest percent identity with the query was selected for STRING analysis. In the “Viewers” section, the “Network” graphic summarizes known or predicted interactions with other proteins, as well as the nature of evidence for each interaction (such as from laboratory experiment, co-expression, or automated textmining).

The database provides any annotations the interacting proteins may have. The display settings did not include automated textmining evidence of interactions (which may be unreliable due to low criteria and lack of hand-curation), had confidence level set to low (0.150 or above; textmining raised confidence scores quite significantly, so this was done to offset the impacts of ignoring it), and with the top 50 first shell-interaction results (so as to have a large pool of potentially significant interactions).

### Subcellular localization

The deep learning model improvement to the Transmembrane Helices Hidden Markov Model (DeepTMHMM v. 1.0.44, located at https://dtu.biolib.com/DeepTMHMM) algorithm was used to detect and predict transmembrane alpha helices and beta barrels within query protein sequences[18]. The FASTA protein sequence was run through the algorithm and the resulting schematics indicated which regions of the protein, if any, are likely transmembrane domains, as well as the probability of this prediction and the most likely topology of the overall protein.

The SignalP v. 6.0 server (https://dtu.biolib.com/SignalP-6) was utilized to predict if query proteins contain signal peptides (SPs). The machine learning model is able to detect all five types of SPs and their associated features, such as cleavage sites, based on information from protein sequences across all domains of life[19]. The protein FASTA sequence was inputted, and the organism was set to eukarya with the default fast mode. The results indicated the likelihood of the presence of any SP, with “Other” meaning no SP was present. It also indicated the likely cleavage site and specific N-H-C regions of the SP.

To determine where a protein of interest is likely targeted, the TargetP v. 2.0 server (located at https://services.healthtech.dtu.dk/services/TargetP-2.0/) was used. TargetP, using deep learning via deep neural networks, recognizes patterns in the N-terminal to predict the presence of mitochondrial transit peptides (mTP), chloroplast transit peptide (cTP), thylakoid luminal transit peptide (lTP), or SPs which typically indicate the protein is targeted to the secretory pathway[20]. The FASTA sequence was pasted, the organism specified as a “plant,” and the long output format was used. The results showed the likelihood of each type of N-terminal presequence, as well as corresponding cleavage sites. A likelihood of “other” indicated that no presequence was detected by the algorithm.

The WoLF PSORT (Protein Subcellular Localization Prediction) tool (located at https://wolfpsort.hgc.jp/) was used to supplement SignalP and TargetP in regards to where proteins of interest localize to. WoLF PSORT determines the query protein’s neighbors based on localization features, and then uses various signals and functional motifs within the neighbor proteins to determine their localization and the query protein’s potential localization[21]. The protein FASTA sequence was entered in the “text area” and the organism was selected as a “plant.” The top nearest neighbors and their localizations were listed, as well as any annotations they may have. In the “Normalized Feature Values” table, the percentile values of each localization feature (such as signals targeting the protein for mitochondria, vacuoles, or the nucleus) was provided for the query protein and its neighbors, with a higher value suggesting a stronger signal for that feature.

Additionally, the functional enrichment analysis resource in STRING was used to determine the subcellular localization of the proteins that interact with the query protein, which may indicate the localization of the query. Found under the “Analysis” tab in STRING, functional enrichment analysis is conducted regarding GO terms present among the output set of proteins. In observing data, the default setting of the “signal” was used to determine enrichment, which is defined as a weighted harmonic mean between the observed/expected ratio (measures how large the enrichment effect is) and the −log(FDR) (false discovery rate, measures how significant enrichment is as a p-value); the analysis looked at was of “cellular compartment” GO terms.

## Results

### *Ab initio* gene prediction and protein alignment

Genes of interest were selected and narrowed down based on the length of the AUGUSTUS-predicted protein sequence (250-750 amino acids) and the relative quality of their BLASTP alignments (preferring the presence of many hits with both query coverage and percent identity over 85%, and E-values on a lower order of magnitude than 10^−5^) and MSA Viewer conservations (more conserved regions and fewer insertions, deletions, or variable regions). The ultimately chosen gene was the 86796-Op.00g065710.m01-00001 gene located on contig tig00000022, comprising bases 16,594,582-16,599,948. The AUGUSTUS gene prediction contained 695 amino acid residues, with 10 exons (Fig 1A). The local alignment search on BLAST indicated high conservation across homologs, with the top 50 results having query coverage of 97-100%, percent identity of 87-95%, and E-values of 0.0 for all. To further assess alignment with other species on MSA Viewer, 8 homologs were hand-chosen (Table 1). Overall, the sequences were conserved fairly well, indicated in red (Fig 1B).

**Table 1.** Top 8 hand-selected homologs to assess alignment. Selection prioritized choosing proteins within the top 50 BLAST hits from different species and with accession codes beginning with XP or NP.

| Description | Organism | Query Cover | E-Value | % Identity | Accession |
| --- | --- | --- | --- | --- | --- |
| Predicted: oxysterol-binding protein-related protein 2B isoform X1 | *Camelina sativa* | 100% | 0.0 | 94.68% | XP_019089498.1 |
| OSBP(oxysterol binding protein)-related protein 2B | *Arabidopsis thaliana* | 86% | 0.0 | 92.85% | NP_001327967.1 |
| Oxysterol-binding protein-related protein 2B isoform X2 | *Capsella rubella* | 100% | 0.0 | 92.23% | XP_023636284.1 |
| Low quality: oxysterol-binding protein-related protein 2B | *Arabidopsis lyrata* subsp. *lyrata* | 100% | 0.0 | 91.80% | XP_020877647.1 |
| Oxysterol-binding protein-related protein 2B isoform X1 | *Eutrema salsugineum* | 100% | 0.0 | 89.41% | XP_024010108.1 |
| Oxysterol-binding protein-related protein 2B isoform X1 | *Brassica rapa* | 100% | 0.0 | 88.84% | XP_009134270.2 |
| Oxysterol-binding protein-related protein 2B isoform X2 | *Brassica napus* | 100% | 0.0 | 88.70% | XP_048606944.1 |
| Oxysterol-binding protein-related protein 2B isoform X1 | *Raphanus sativus* | 100% | 0.0 | 88.03% | XP_018476566.1 |

**Fig 1.**
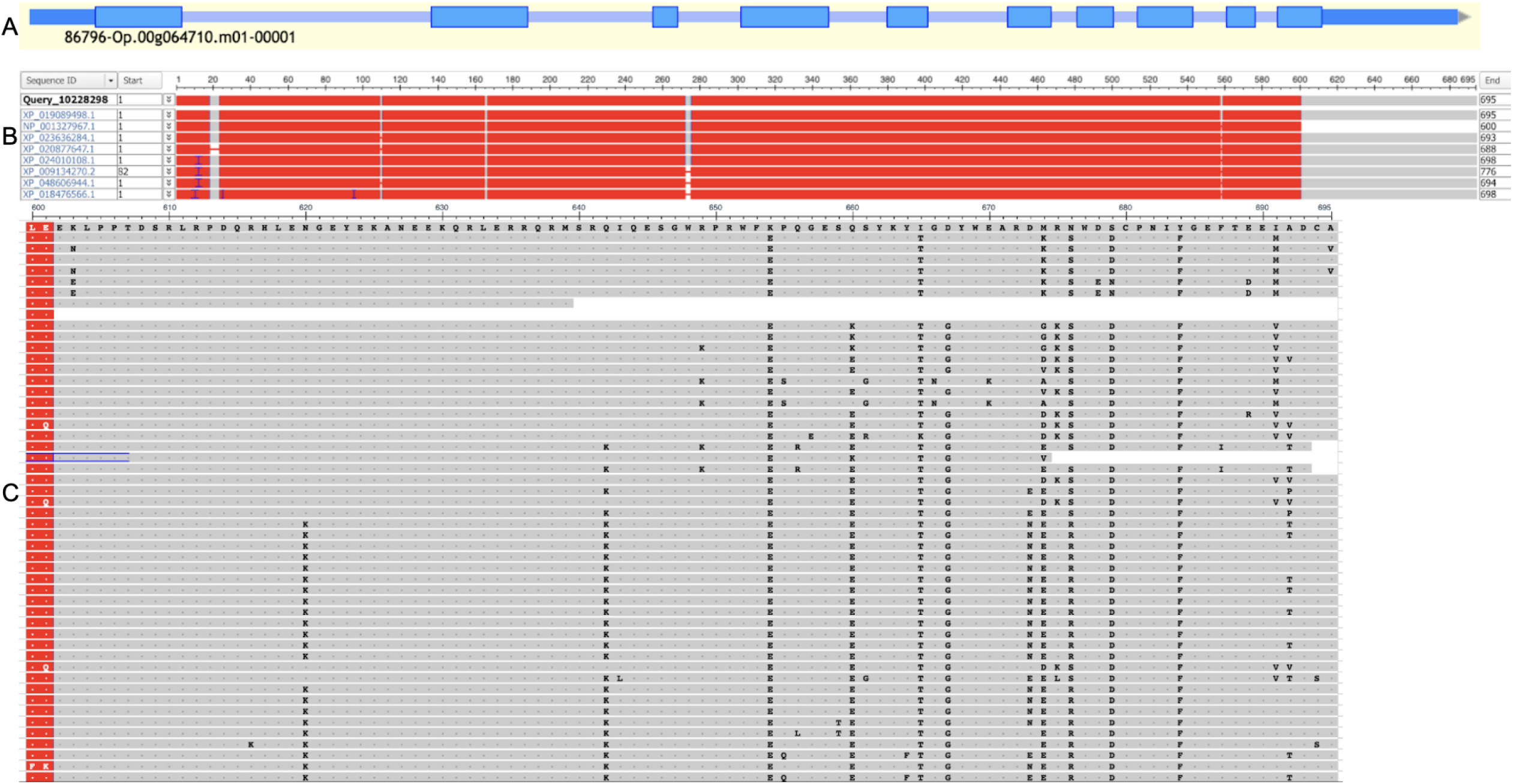
Homology-based gene annotation of oxysterol-binding protein-related protein. (A) Augustus track of gene 86796-Op.00g064710.m01-00001 in tig00000022 of *O. pumila*, viewed in GeneSAS. (B) MSA Viewer of protein sequence. In red is conserved regions, with areas of low conservation in gray. Thin red lines indicate a deletion in comparison with the query, and a blue bracket indicates an insertion in comparison with the query. (C) Zoomed-in view of C-terminus, residues 600-695, of the homologs’ sequences. Dots indicate the same amino acid as the query protein, and amino acid 1-letter symbols indicate residues that differ from the query.

Regions with deletions or insertions were only affected in some of the homologs while the others had the same sequence as the query protein, indicating that there was likely no error in the AUGUSTS prediction of those regions and that it was likely due to natural variation. Notably, the C-terminus of the protein (amino acids 602-695) is not conserved among homologs, with this region even being deleted in the protein from *A. thaliana*. When compared with the top 50 BLAST hits, this region seems to be largely variable among homologs (Fig 1C). Certain residues, such as Lys654 and Tyr684 in the query protein, are fully conserved as a different residue in all the other homologs (e.g. Glu654 and Phe684), probably indicating either a sequencing error or a point mutation in the sequenced individual. However, because these differences are within a largely variable segment and they affect only some individual amino acids, we did not manually edit the predicted *O. pumila* protein.Similarly, due to the lack of any large differences (i.e. conserved insertional or deletional differences between the *O. pumila* query protein and its homologs), the AUGUSTUS prediction for this gene was not altered and was concluded to be the most refined model.

### Structural analysis

After conservation was analyzed and the AUGUSTUS prediction was determined to be an accurate model of the 86796-Op.00g065710.m01-0000 gene, we sought to determine the structure of the protein. PSIPRED indicated that the secondary structure of the protein includes multiple alpha helices and beta strands, connected by various coils (Fig 2A). AlphaFold visualized the three-dimensional structure of the protein, with three main confident domains (Fig 2B). The domain starting at the approximately 310th amino acid until the end of the peptide has a fairly large confidence in its structure (mainly blue coloring, meaning higher plDDTs), with very low expected position errors (Fig 2C). The coil and alpha helix consisting of residues 160-240 also have low expected position errors, but lower local confidence (indicated by more light blue and yellow coloring in the structure).

**Figure 2.**
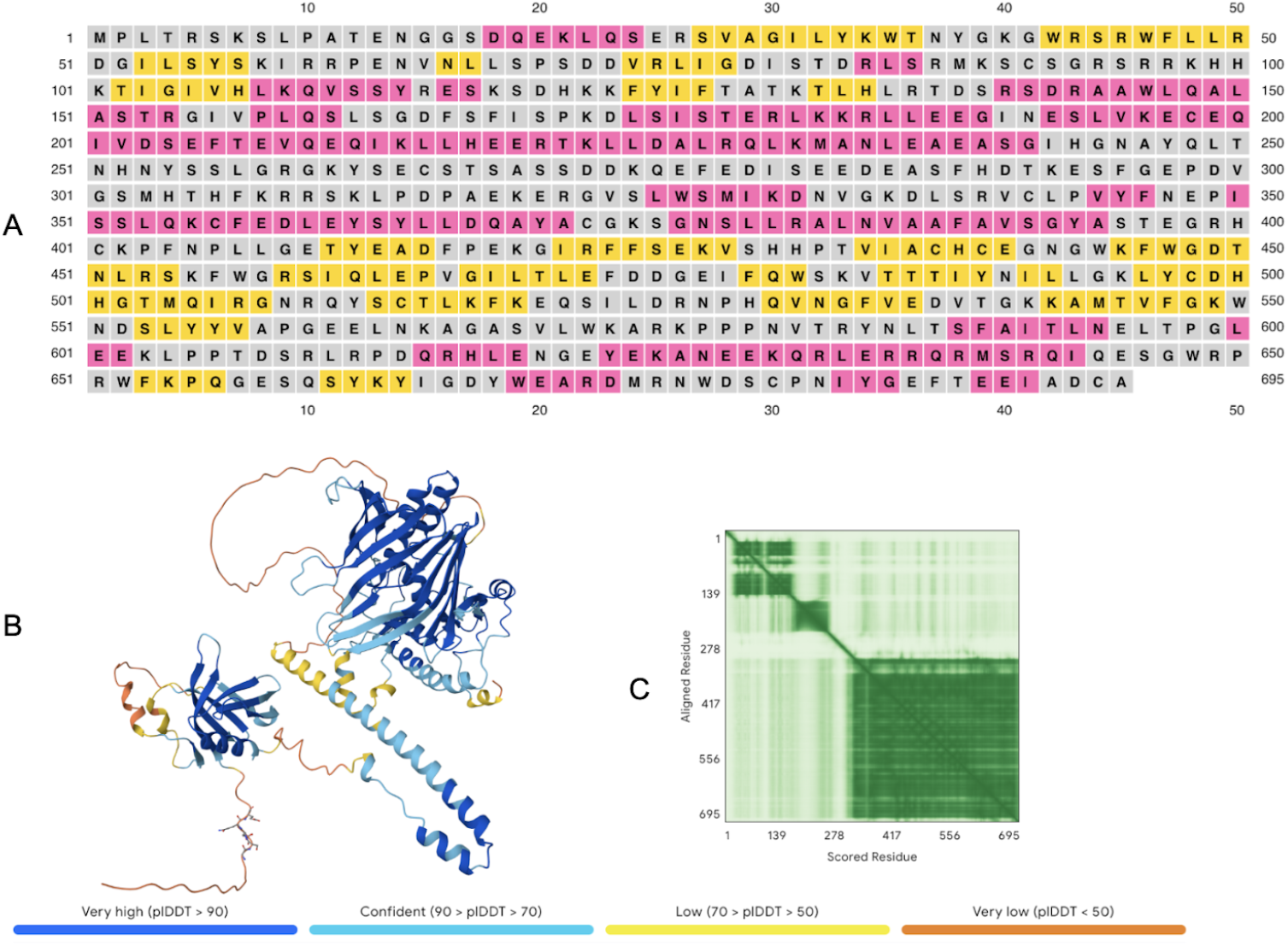
Structural prediction of the oxysterol-binding protein-related protein. (A) PSIPRED secondary structure prediction. Yellow indicates a beta strand, pink an alpha helix, and gray a coil. (B) AlphaFold 3D structure prediction and plDDT color legend. plDDT scores indicate confidence of local position and structure. The pTM score measuring the confidence of the overall structure prediction was 0.6. (C) Expected position error graph (units: Angstroms) of each residue in relation to each other residue. Darker green indicates lower error.

The domain comprising residues 20-140 has more segments that are less confident in local structure and position, but also has a region of higher confidence (dark blue coloring). Each of these domains is connected by coils of low local confidence (colored in orange). The pTM score measuring the overall confidence of the entire structure is 0.6, indicating a fairly high chance that this displayed structure is similar to the true protein structure. We tested whether or not the protein binds other identical peptides to form a dimer, trimer, or other polymer of multiple identical subunits; the pTM scores decreased with increasing number of subunits, which we took as an indication that this peptide does not bind with other identical copies to form a larger protein (S1 Fig).

### Functional analysis

The function of the protein was analyzed using InterPro. InterPro predicted the protein to be oxysterol-binding protein-related (Fig 3A) — meaning it likely binds a variety of oxysterols to assist with or regulate sterol synthesis[22,23] — which agreed with the top BLAST homologs (Table 1). The protein contains an oxysterol binding domain (IPR037239) in the C-terminal half, starting at about the 320th amino acid, and a pleckstrin homology(PH)-like domain (IPR001849) covering residues 26-157, which typically serve as simple targeting domains that bind lipids[24]. These functional domains align with the high-confidence structural domains predicted by AlphaFold, with the PH-like domain correlating with the first structural domain and the oxysterol binding domain correlating with the third, most confident structural domain. There are also regions of InterPro-

**Figure 3.**
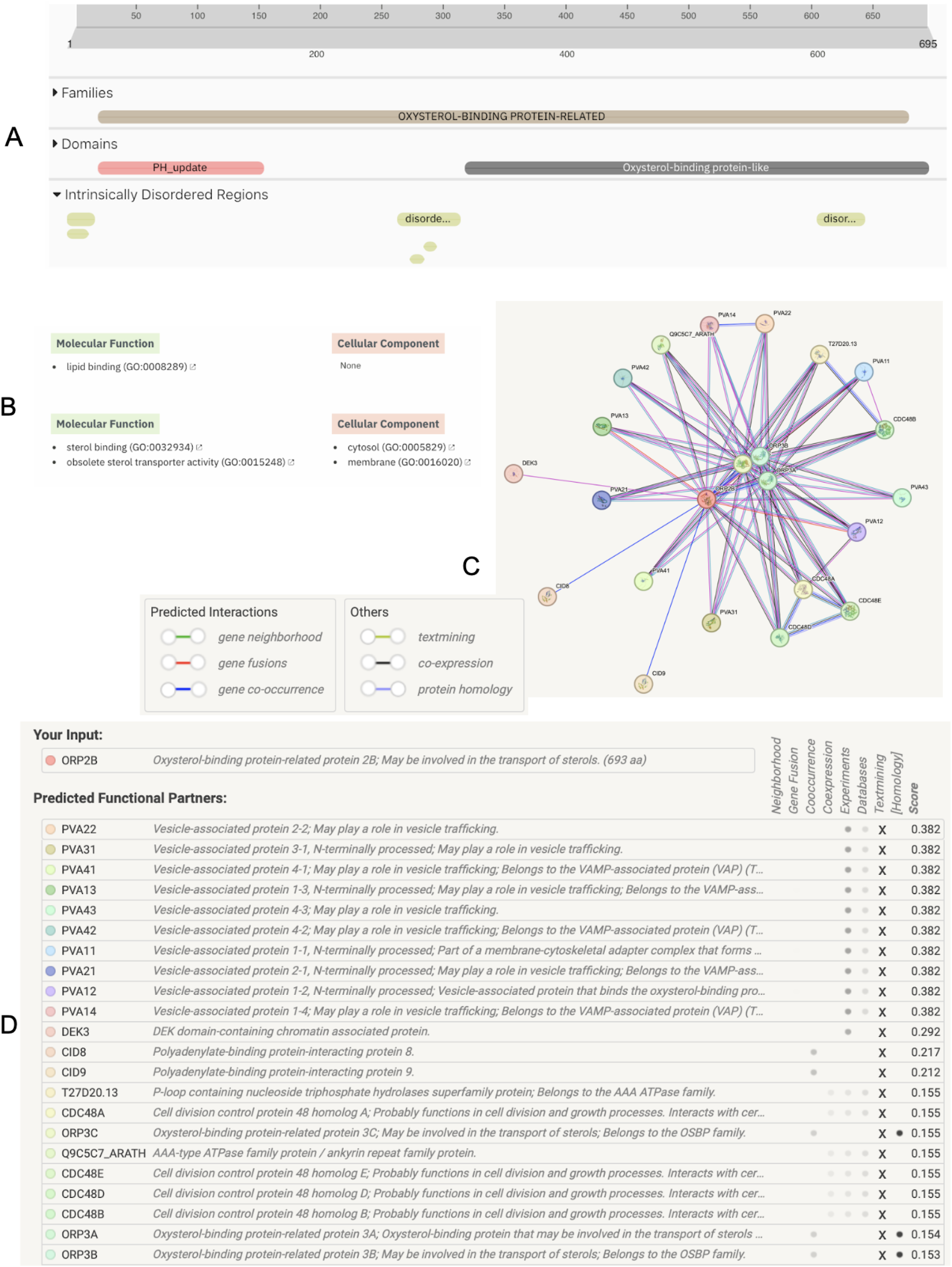
Functional analysis of the oxysterol-binding protein-related protein. (A) InterPro family, domain, and intrinsic disorder prediction map. (B) All InterPro-predicted GO terms for the OSBP-related protein. (C) STRING prediction of proteins interacting with ORP2B and color legend of interaction/evidence types. (D) Gene annotations of these interacting proteins and the nature of evidence of interactions.

I predicted intrinsic disorder at residues 1-22, 265-314, and 600-637, which align with some of the coils of low structural prediction confidence in between the more confident domains found in AlphaFold. InterPro provided gene ontology (GO) terms relating to the protein’s predicted molecular function and cellular compartment, particularly sterol binding (GO:0032934) in the cytosol (GO:0005829) and/or in membranes (GO:0016020) (Fig 3B).

We further assessed this oxysterol-binding protein(OSBP)-related protein’s function using STRING to determine any direct or indirect interactions it may be involved in. Searching the protein sequence in the database with the organism set to *A. thaliana* matched it most directly with OSBP-related protein (ORP) 2B,which is likely involved in the transport of sterols. This prediction agreed with the outputs of both BLAST (Table 1) and InterPro (Fig 3A). The STRING database predicts ORP2B to interact with various proteins of note (Fig 3C-D). Among these are vesicle-associated proteins involved in vesicle trafficking like PVA11, which is part of a membrane-cytoskeletal complex that forms a bridge between the endoplasmic reticulum (ER) and the plasma membrane. Other predicted interacting partners were cell division control proteins like CDC48A, which are involved in cell growth and/or division via interactions with certain SNARE (soluble N-ethylmaleimide-sensitive factor attachment protein receptor) proteins that participate in homotypic fusion of vesicles. Lastly, ORP2B demonstrates some level of co-occurrence with ORPs 3A-C, which are involved in the transport of sterols from the ER to the Golgi apparatus (Fig 3D).

### Transmembrane domains and subcellular localization

After predicting the structure and function of the protein product of the 86796-Op.00g065710.m01-00001 gene in *O. pumila*, which was found to be ORP2B, we then sought to determine the protein’s subcellular localization. DeepTmHMM predicted a 100% likelihood that ORP2B has no transmembrane domains (S2 Fig). Similarly, both SignalP (S3 Fig) and TargetP (S4 Fig) predicted almost 100% likelihood of no signal peptides or other target sequences being present. Interestingly, WoLF PSORT predicted ORP2B to potentially localize in the nucleus based on a large majority of the localization feature neighbors and on the query protein’s localization features themselves, though the score for nuclear localization of ORP2B was only in the 61st percentile (Fig 4A). In contrast, the GO functional enrichment from the STRING database demonstrated that a significant portion of the proteins that ORP2B interacts and likely shares a function with localize near the ER (Fig 4B).

**Figure 4.**
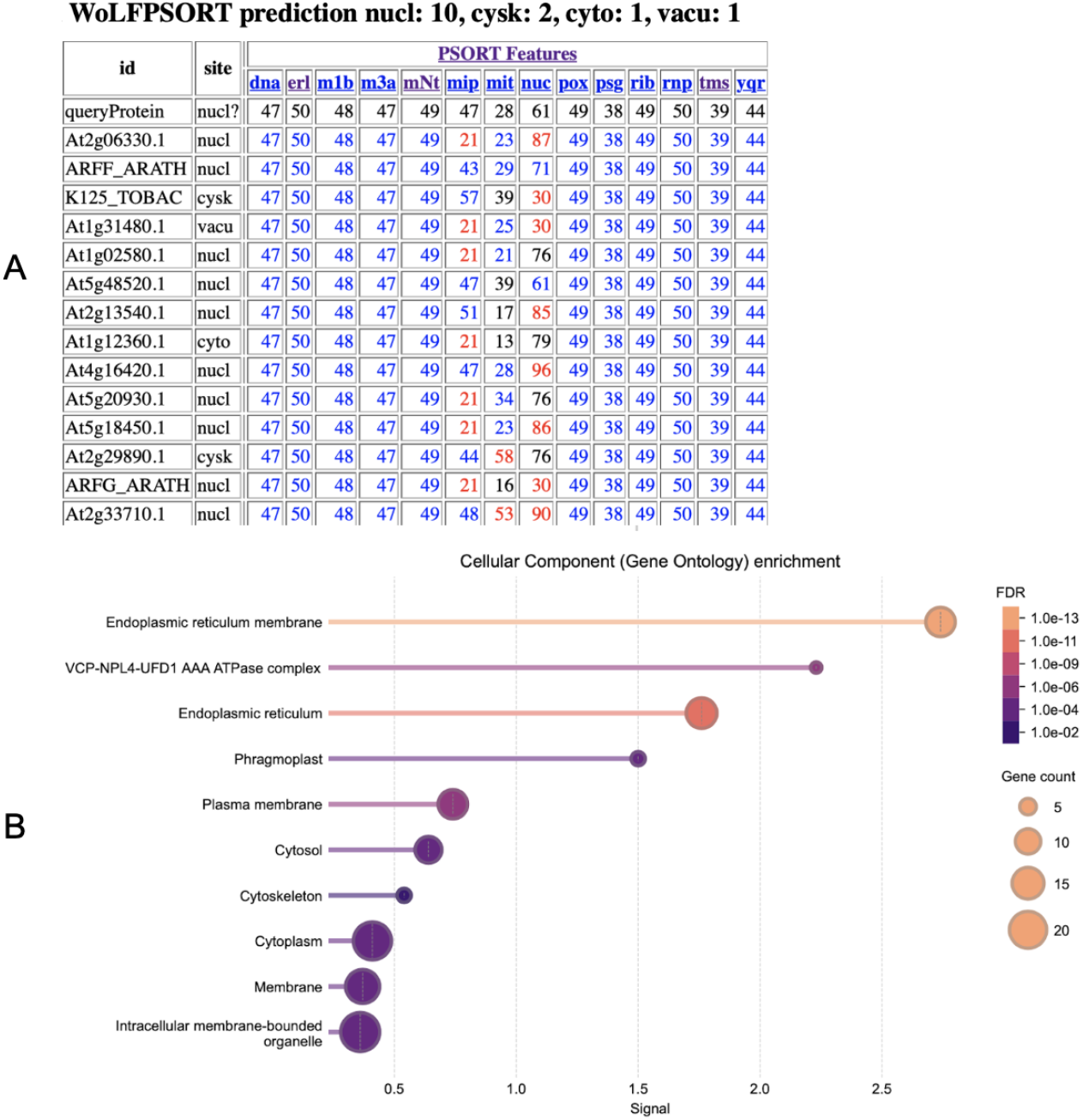
Subcellular localization predictions for the oxysterol-binding protein-related protein 2B. (A) WoLF PSORT predictions based on neighbors (top) and percentile value for localization features of the query protein and its neighbors (bottom), dna and nuc = nuclear proteins; erl = ER lumen; mlb, m3a, mNt, and tms = membrane proteins; mip and mit = mitochondria; pox = peroxisome; psg = signal peptide; rib = ribosomal proteins; mp = RNA-binding proteins; yqr = endocytotic membrane proteins. (B) Functional enrichment visualization of STRING output proteins’ cellular component GO terms. The VCP-NPL4-UFD1 AAA ATPase complex is involved in essential ER processes and activity.

## Discussion

This study manually annotated and explored the conservation, structure, function, and localization of the AUGUSTUS-predicted 86796-Op.00g065710.m01-00001 gene located on contig tig00000022 of the *O. pumila* genome, comprising bases 16,594,582-16,599,948. We validated the AUGUSTUS gene model based on BLAST and MSA Viewer alignments with homologs (Fig 1B). We concluded, based on BLAST (Fig 1B), InterPro (Fig 3A), and STRING (Fig 3D), that this gene encodes the oxysterol-binding protein-related protein(ORP)-2B, which is conserved fairly well across related plant species. Structurally, ORP2B’s PH domain in the first approximately 150 amino acid residues is a feature shared with other ORPs in *A. thaliana*, such as ORPs 1A-D and ORP2A, although it is missing in others, such as ORPs 3A-C and 4A-C[25]. Regarding the protein’s localization within the cell, we found conflicting information. SignalP and TargetP agreed that there are no signal peptides or target presequences within the peptide, and the analysis by DeepTMHMM indicates the protein does not span any membranes via a transmembrane domain. However, WoLF PSORT predicted, based largely on the protein’s “neighbors,” that ORP2B localizes to the nucleus (Fig 4A).

Meanwhile, the functional enrichment analysis from STRING indicated that a large portion of the proteins that may interact with ORP2B are localized to the ER (Fig 4B). Previous research has found that ORP2A, which is similar in sequence to ORP2B, functions primarily on the ER membrane[25,26]. We therefore concluded that it is likely ORP2B localizes to the ER as indicated by the STRING database and previous studies, although further research should be completed to provide more substantive and direct evidence.

While ORP2B itself is not well-studied, previous research has been conducted regarding OBSPs and ORPs in general, as well as other specific ORPs from *A. thaliana*. ORP2A has been found to mediate ER-autophagosomal contacts through the redistribution of phosphatidylinositol 3-phosphate (PI3P) in the ER membrane, and ORP2A knockdowns in *A. thaliana* seedlings not only have impaired autophagy but also stunted growth[26]. This indicates an essential role of ORP2A, and therefore likely ORP2B, in cell growth and proliferation. In human cells, ORP2 has similarly been found to regulate cholesterol transport to the plasma membrane through the exchange of phosphatidylinositol 4,5-bisphosphate (PI(4,5)P_2_)[27]. ORP4 in mammals also binds phosphatidylinositol 4-phosphate (PI4P), and this activity promotes cell survival and division[28]. In zebrafish, ORP2 plays an important role in fat cell differentiation and growth[29]. Additionally, ORPs and the VAPs (vesicle-associate proteins) they bind to — including ORP2B, as indicated by the STRING results (Fig 3D) — have been shown to be co-opted by viruses to assist viral replication in plants and yeast[30], as well as to promote and accelerate cancer cell growth, proliferation, and migration in mammals[31]. It is therefore possible, and perhaps likely, that ORP2B binds to a form of PIP in its function of sterol transportation, and that this function is vastly important to cell survival and division. This contribution to cell proliferation by ORP2B is also supported by the STRING database’s indication of interactions and shared function with cell division control proteins (Fig 3D).

Through our analysis, we have concluded that this *ORP2B* gene is important for cell survival, growth, and division via its activity in sterol transportation to or from the ER. We have contributed to the functional annotation of the markedly understudied *O. pumila* genome, which is an understated and potentially significant source of a novel backdrop and natural variation for various fields of scientific research. In fact, the gene analyzed in this study may be a contributor to this promising characteristic of its species: in *A. thaliana*, ORP2B is upregulated in conditions of high light and drought[25], conditions which *O. pumila* is known to have adapted to and in which *O. pumila* has diverged from the model system.

## Acknowledgements

Thank you to Dr. Lukasz Salwinski and Ye Wang for their instruction and guidance.

## Supporting Information

**S1 Fig.**
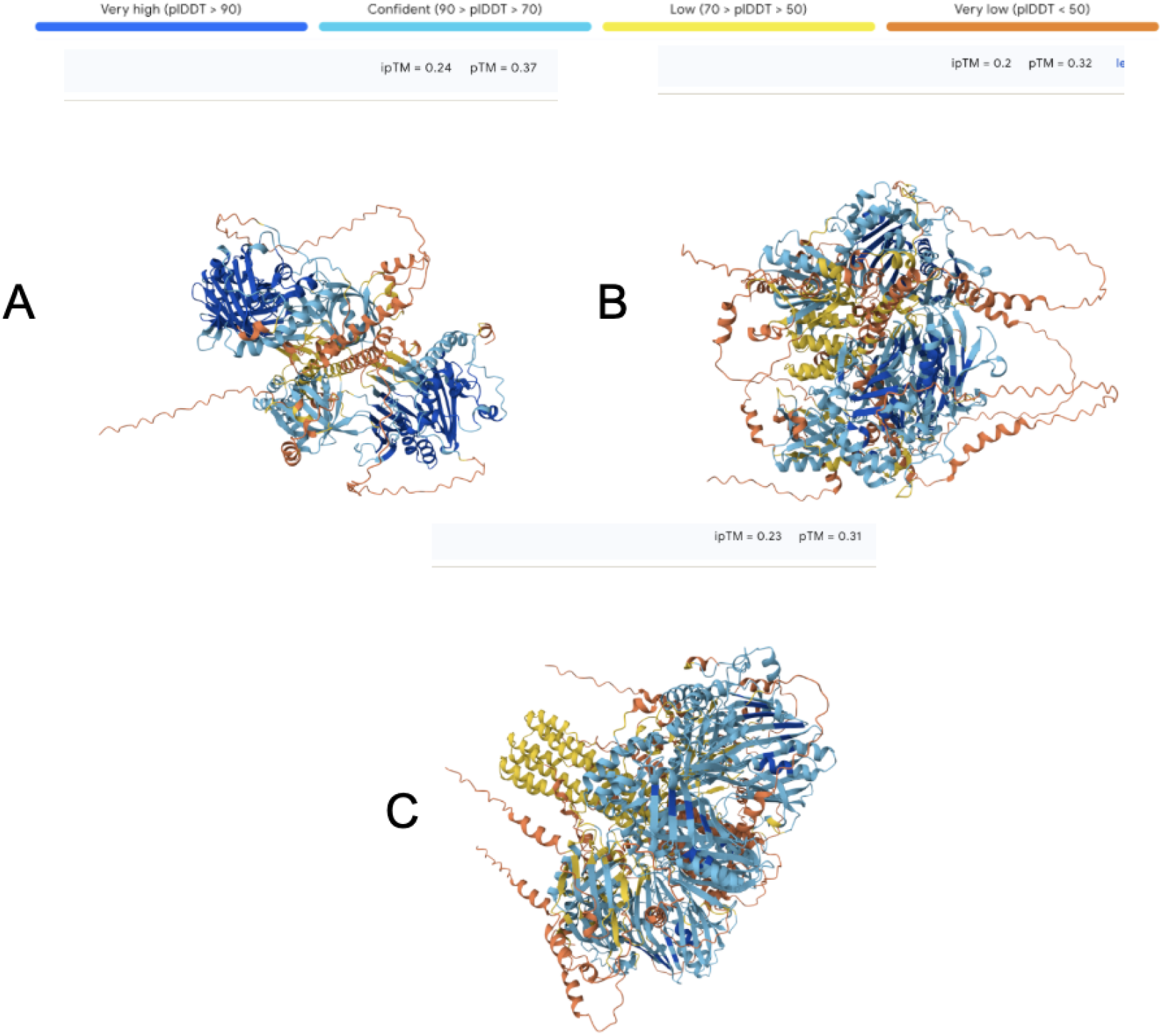
AlphaFold predictions of the quaternary’ structures of a dimer (A), trimer (B), and tetramer (C) of identical copies of ORP2B, and their associated pTM scores and plDDT score coloring. The plDDT scoring indicates local structural and positional confidence. A pTM score (measures overall structural confidence) greater than 0.5 indicates the likelihood that the predicted structure is similar to the natural structure.

**S2 Fig.**
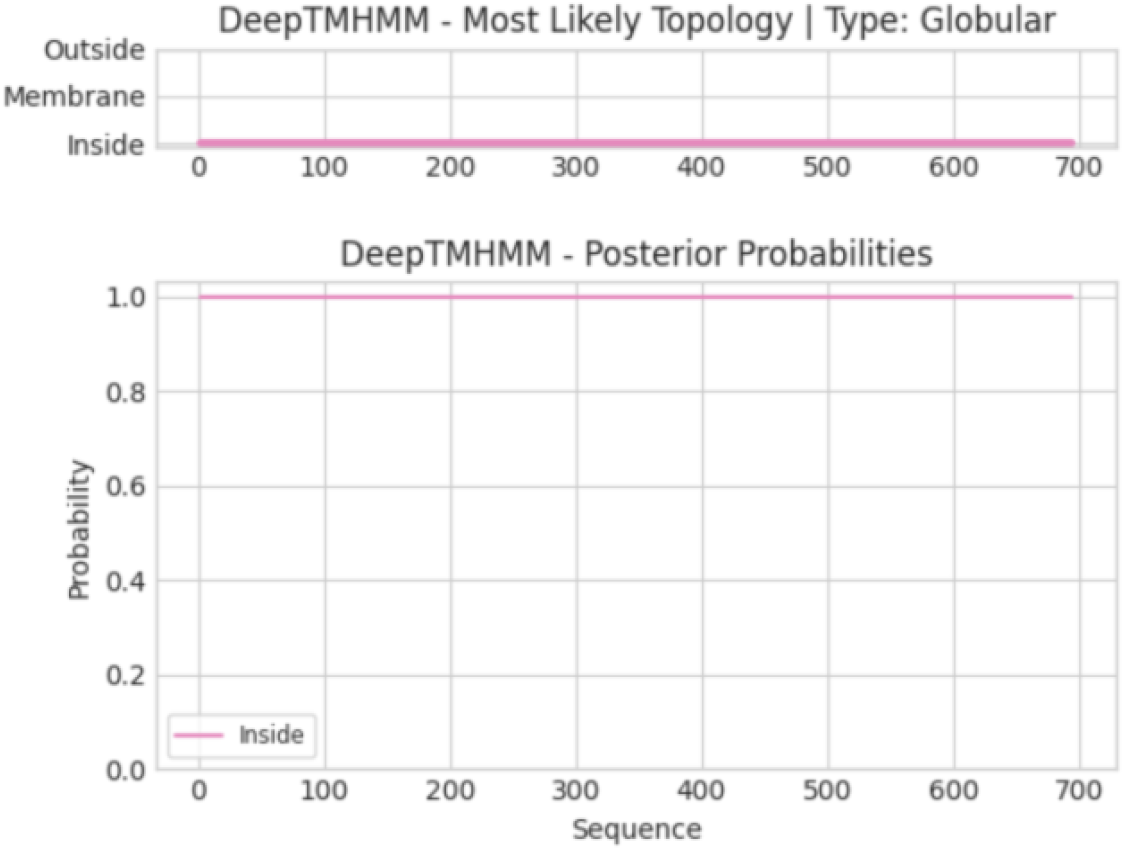
DeepTMHMM prediction of the likely topology of the protein ORP2B.

**S3 Fig.**
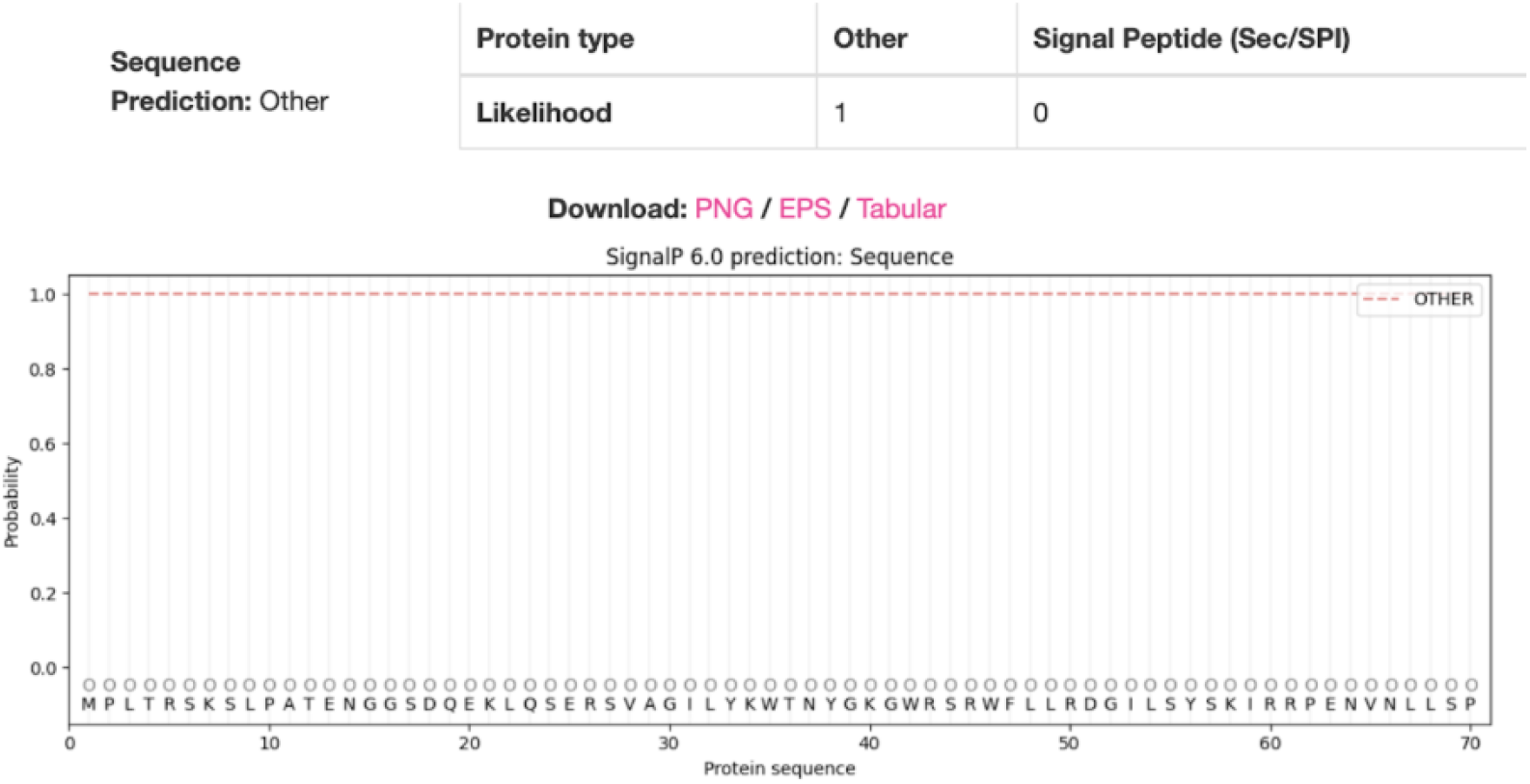
SignalP prediction of the likelihood of signal peptides present in the sequence of ORP2B.

**S4 Fig.**
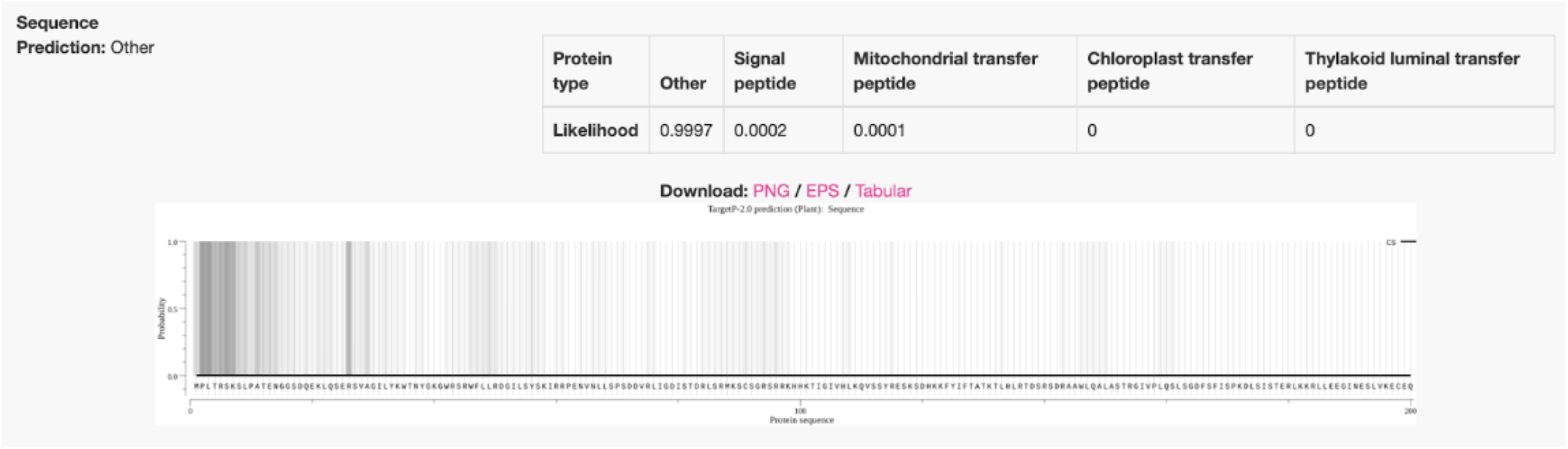
TargetP prediction of the likelihood of various signal peptides of target presequences in the sequence of ORP2B.

## Notes

### Competing Interest Statement

The authors have declared no competing interest.

